# Ocular disease mechanisms elucidated by genetics of human fetal retinal pigment epithelium gene expression

**DOI:** 10.1101/446799

**Authors:** Boxiang Liu, Melissa A. Calton, Nathan S. Abell, Gillie Benchorin, Michael J. Gloudemans, Ming Chen, Jane Hu, Xin Li, Brunilda Balliu, Dean Bok, Stephen B. Montgomery, Douglas Vollrath

**Affiliations:** Department of Biology, Stanford University, Stanford, CA 94305, USA; Department of Genetics, Stanford University School of Medicine, Stanford, CA 94305, USA; Program in Biomedical Informatics, Stanford University School of Medicine, Stanford, CA; Department of Ophthalmology, Jules Stein Eye Institute, UCLA, Los Angeles, CA; Department of Pathology, Stanford University School of Medicine, Stanford, CA 94305, USA 94305, USA

**Author notes:** equal contributions (Boxiang Liu, Melissa A. Calton).

## Abstract

The eye is an intricate organ with limited representation in large-scale functional genomics datasets. The retinal pigment epithelium (RPE) serves vital roles in ocular development and retinal homeostasis. We interrogated the genetics of gene expression of cultured human fetal RPE (fRPE) cells under two metabolic conditions. Genes with disproportionately high fRPE expression are enriched for genes related to inherited ocular diseases. Variants near these fRPE-selective genes explain a larger fraction of risk for both age-related macular degeneration (AMD) and myopia than variants near genes enriched in 53 other human tissues. Increased mitochondrial oxidation of glutamine by fRPE promoted expression of lipid synthesis genes implicated in AMD. Expression and splice quantitative trait loci (e/sQTL) analysis revealed shared and metabolic condition-specific loci of each type and several eQTL not previously described in any tissue. Fine mapping of fRPE e/sQTL across AMD and myopia genome-wide association data suggests new candidate genes, and mechanisms by which the same common variant of *RDH5* contributes to both increased AMD risk and decreased myopia risk. Our study highlights the unique transcriptomic characteristics of fRPE and provides a resource to connect e/sQTL in a critical ocular cell type to monogenic and complex eye disorders.

---

The importance of vision to humans and the accessibility of the eye to examination have motivated the characterization of more than one thousand genetic conditions involving ocular phenotypes ^1^. Among these, numerous monogenic diseases exhibit significant inter- and intra-familial phenotypic variability ^2-7^. Imbalance in allelic expression of a handful of causative genes has been documented ^8^, but common genetic variants responsible for such effects remain to be discovered.

Complementing our knowledge of numerous monogenic ocular disorders, recent genome-wide association studies (GWAS) ^9^ have identified hundreds of independent loci associated with polygenic ocular phenotypes such as age-related macular degeneration (AMD), the leading cause of blindness in elderly individuals in developed countries ^10,11^, and myopia, the most common type of refractive error worldwide and an increasingly common cause of blindness ^12-14^. Despite the rapid success of GWAS in mapping novel ocular disease susceptibility loci, the functional mechanisms underlying these associations are often obscure.

Connecting changes in molecular functions such as gene expression and splicing with specific GWAS genomic variants has aided the elucidation of functional mechanisms. Non-coding variants account for a preponderance of the most significant GWAS loci ^15,16^, and most expression quantitative trait loci (eQTL) map to non-coding variants ^17^. Thousands of eQTL have been found in a variety of human tissues ^18^, but ocular cell-types are underrepresented among eQTL maps across diverse tissues, and no systematic search for eQTL has so far been described for any cell type in the human eye.

The retinal pigment epithelium (RPE) is critical for eye development ^19^ and for an array of homeostatic functions essential for photoreceptors ^20^. Variants of RPE-expressed genes have been associated with both monogenic and polygenic ocular phenotypes, including AMD and myopia. We recently implicated an eQTL associated with an RPE-expressed gene as modulating the severity of inherited photoreceptor degeneration in mice ^21^. To investigate the potential effects of genetically encoded common variation in human RPE gene expression, we set out to identify eQTL in cultured human fetal RPE (fRPE) cells and assess the implications for ocular disease.

## Results

### The transcriptome of human fRPE cells

We studied 23 primary human fRPE lines (Table S3), all generated by the same method in a single laboratory ^22^ and cultured for at least 10 weeks under conditions that promote a differentiated phenotype ^23^. DNA from each line was genotyped at 2,372,784 variants. Additional variants were imputed and phased using Beagle v4.1 ^24^ against 1000 Genomes Phase 3 ^25^ for a total of ~13 million variants after filtering and quality control (see **Methods**).

Our goal was to identify RPE eQTL relevant to the tissue’s role in both developmental and chronic eye diseases. The balance between glycolytic and oxidative cellular energy metabolism changes during development and differentiation ^26^, and loss of RPE mitochondrial oxidative phosphorylation capacity may contribute to the pathogenesis of AMD ^27^. We therefore obtained transcriptional profiles of each fRPE line cultured in medium that favors glycolysis (glucose plus glutamine) and in medium that promotes oxidative phosphorylation (galactose plus glutamine) ^28^. We performed 75-base paired-end sequencing to a median depth of 52.7 million reads (interquartile range: 45.5 to 60.1 million reads) using a paired sample design to minimize batch effects. To determine the relationship between primary fRPE and other tissues, we visualized fRPE against 53 tissues from the GTEx Project v7 ^18^. The fRPE samples formed a distinct cluster situated between heart and skeletal muscle and brain (Fig. 1A), tissues that, like the RPE, are metabolically active and capable of robust oxidative phosphorylation.

**Figure 1.**
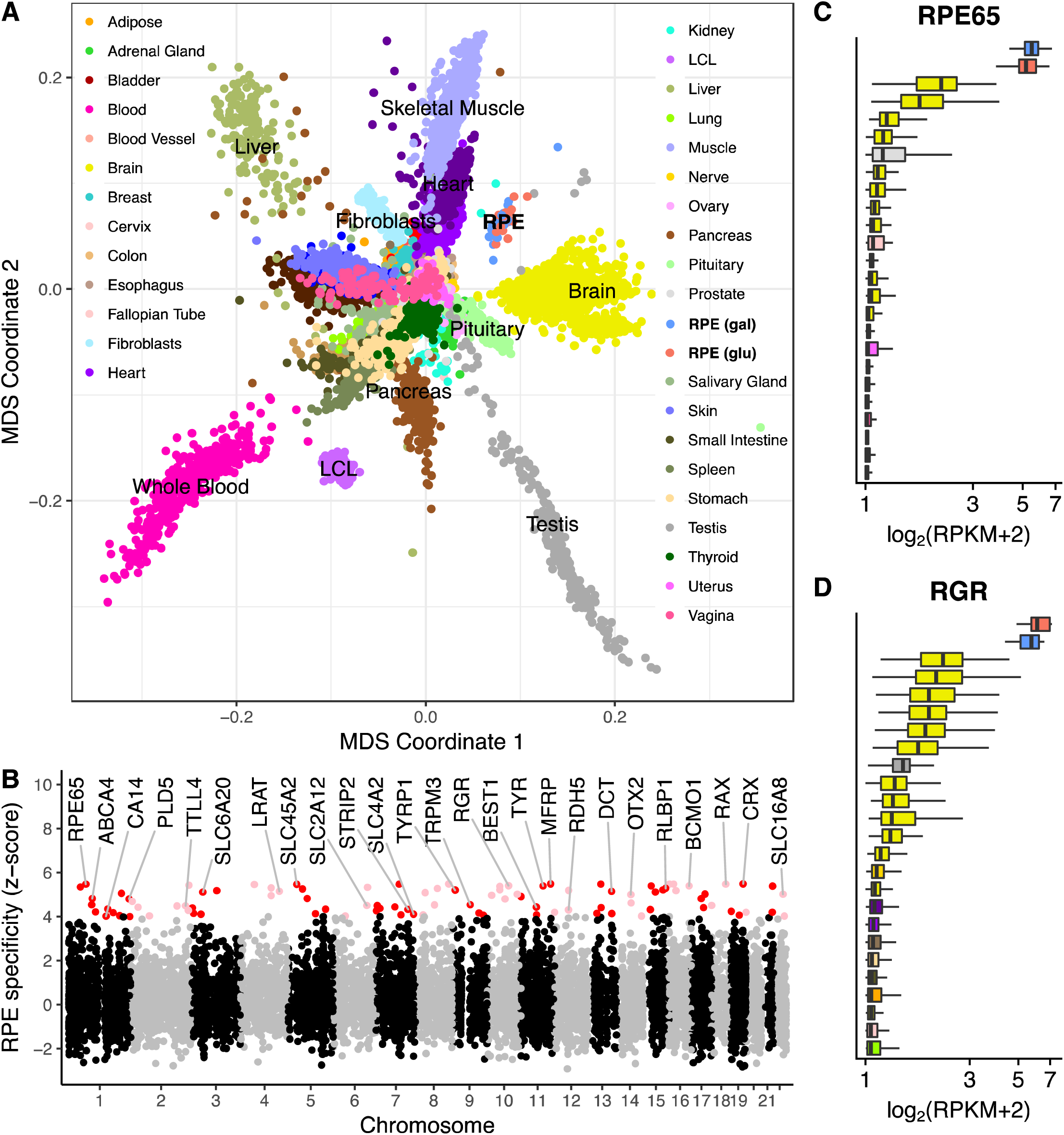
Characteristics of the fRPE transcriptome. (A) Multidimensional scaling against GTEx tissues locates fRPE near heart, skeletal muscle and brain samples. (B) A subset of the fRPE-selective gene set defined by z-score > 4 is shown including RPE signature genes such as *RPE65* and new genes such as *TYR*. Red/pink dots indicate fRPE-selective genes with z-score > 4 in both glucose and galactose conditions. (C-D) Two examples of the expression levels of fRPE-selective genes in various GTEx tissues. Only the top 25 tissues are plotted for visual clarity. For (A), (C) and (D), red indicates fRPE glucose condition and blue indicates fRPE galactose condition.

To identify genes with disproportionately high levels of expression in the fRPE, we compared the median RPKMs of fRPE genes against GTEx tissues. We defined fRPE-selective genes as those with median expression at least four standard deviations above the mean (see **Methods**). Under this definition, we found 100 protein-coding genes and 30 long non-coding RNAs (lncRNAs) to be fRPE-selective (Fig. 1B and Table S5). Multiple previously defined RPE ‘signature’ genes ^29-31^ are present in our list including *RPE65* (Fig. 1C) and *RGR* (Fig. 1D). Using this set of genes, we performed Gene Set Enrichment Analysis (GSEA) ^32^ against 5,917 gene ontology (GO) annotations ^33^. The two gene sets most enriched with fRPE-selective genes were *pigment granule* and *sensory perception of light stimulus* (FDR < 1×10^−3^), consistent with the capacity of fRPE to produce melanin and the tissue’s essential role in the visual cycle. Other highly enriched pathways relate to protein synthesis and secretion, eye morphogenesis, and cellular metabolism including oxidative phosphorylation. Table S6 lists the 29 GO pathways enriched using a conservative FWER < 0.05.

### Transcriptomic differences across two metabolic conditions

To gain insight into the response of fRPE cells to altered energy metabolism, we compared gene expression between the two culture conditions using DESeq2 ^34^, correcting for age, sex, ancestry, RIN and batch (see **Methods**). A total of 837 genes showed evidence of significant differential expression (FDR < 1×10^−3^, Fig. 2A and Table S7). Notably, three of the top five differentially expressed genes are involved in lipid metabolism (*SCD*, *INSIG1*, and *HMGCS2* in order). *SCD* codes for a key enzyme in fatty acid metabolism ^35^, and its expression in RPE is regulated by retinoic acid ^36^. *INSIG1* encodes an insulin-induced protein that regulates cellular cholesterol concentration ^37^. *HMGCS2* encodes a mitochondrial enzyme that catalyzes the first step of ketogenesis ^38^, and this enzyme plays a key role in phagocytosis-dependent ketogenesis in fRPE ^39^. To understand the broader impact induced by changes in energy metabolism, we performed pathway enrichment analysis using GSEA ^32^ and found that the top two upregulated pathways in galactose medium are cholesterol homeostasis and mTORC1 signaling (FDR < 1×10^−4^, Fig. 2B). Consistent with the cholesterol finding, forcing cells to rely primarily on oxidation of glutamine for ATP generation increases expression of a suite of genes that promotes lipid synthesis and import (Fig. 2C).

**Figure 2.**
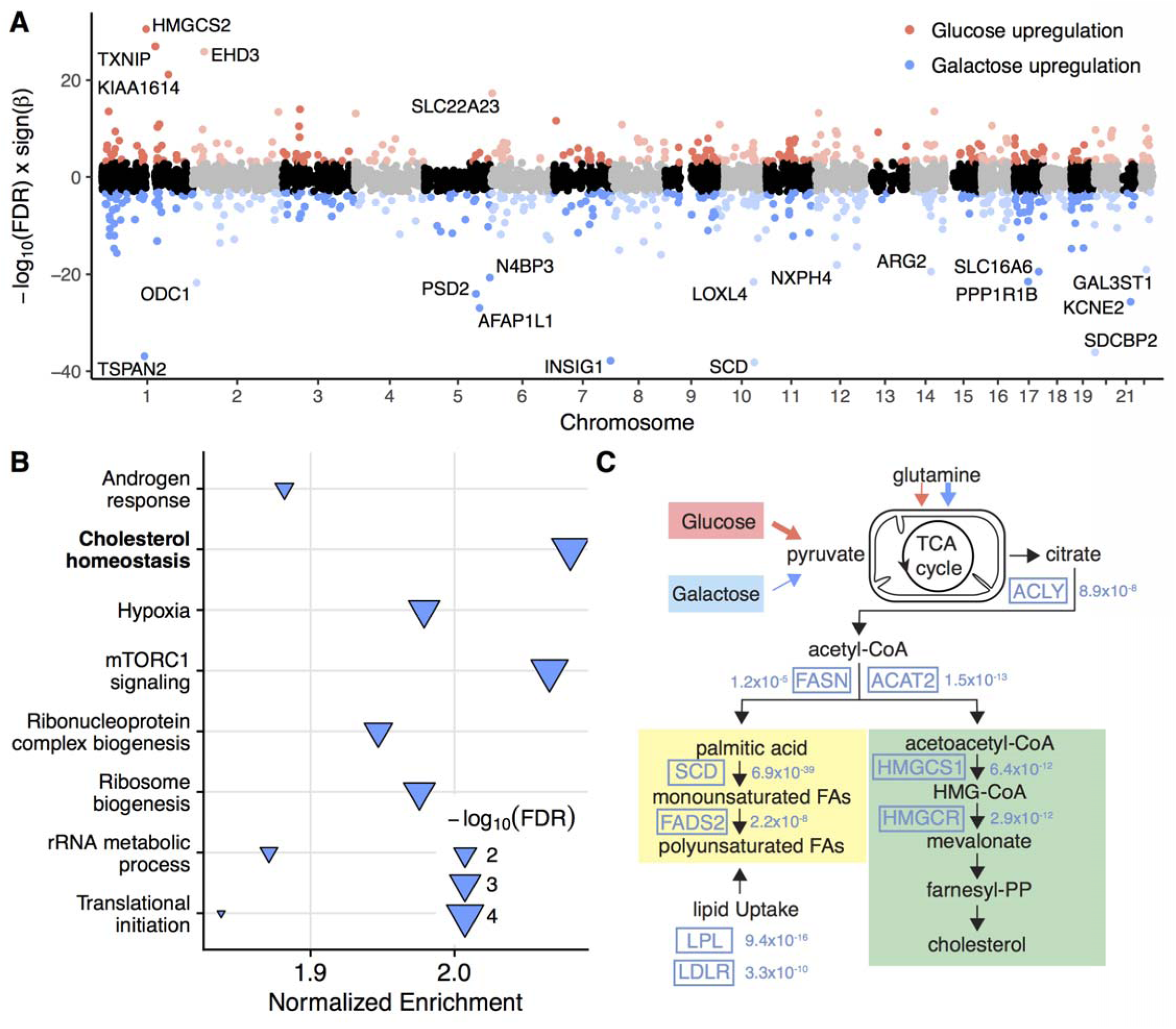
Differential expression across two metabolic conditions. (A) Transcriptome-wide differential expression patterns: red indicates upregulated in glucose, blue indicates upregulated in galactose. (B) G Gene set enrichment analysis of differentially expressed genes. The pathway most enriched is cholesterol homeostasis (upregulated in galactose condition). (C) Key genes involved in cholesterol biosynthesis an and import are upregulated in response to the increased oxidation of glutamine that occurs in the galactose condition. Estimated FDR values are shown next to the gene names.

### fRPE-selective genes are enriched in genetic ocular diseases

Disease-associated genes can have elevated expression levels in effector tissues ^40^. To determine whether ocular disease genes have elevated expression levels in fRPE, we used a manually curated list of 257 ocular disease related genes ^41^ that includes genes for inherited retinal disorders (n = 238), glaucoma (n = 8) and other risk genes (n = 11, Table S2). Compared to all other protein-coding genes, ocular disease-related genes are expressed at higher levels in fRPE (two-sided t-test p-value: 1.6×10^−10^). Further, ocular disease gene expression demonstrated a larger elevation in fRPE than GTEx tissues (Fig. 3A), suggesting fRPE as a model system for a number of eye diseases. As a control, we repeated the analysis for epilepsy genes (n = 189) and observed elevated expression levels in brain tissues as we expected (Fig. S16).

**Figure 3.**
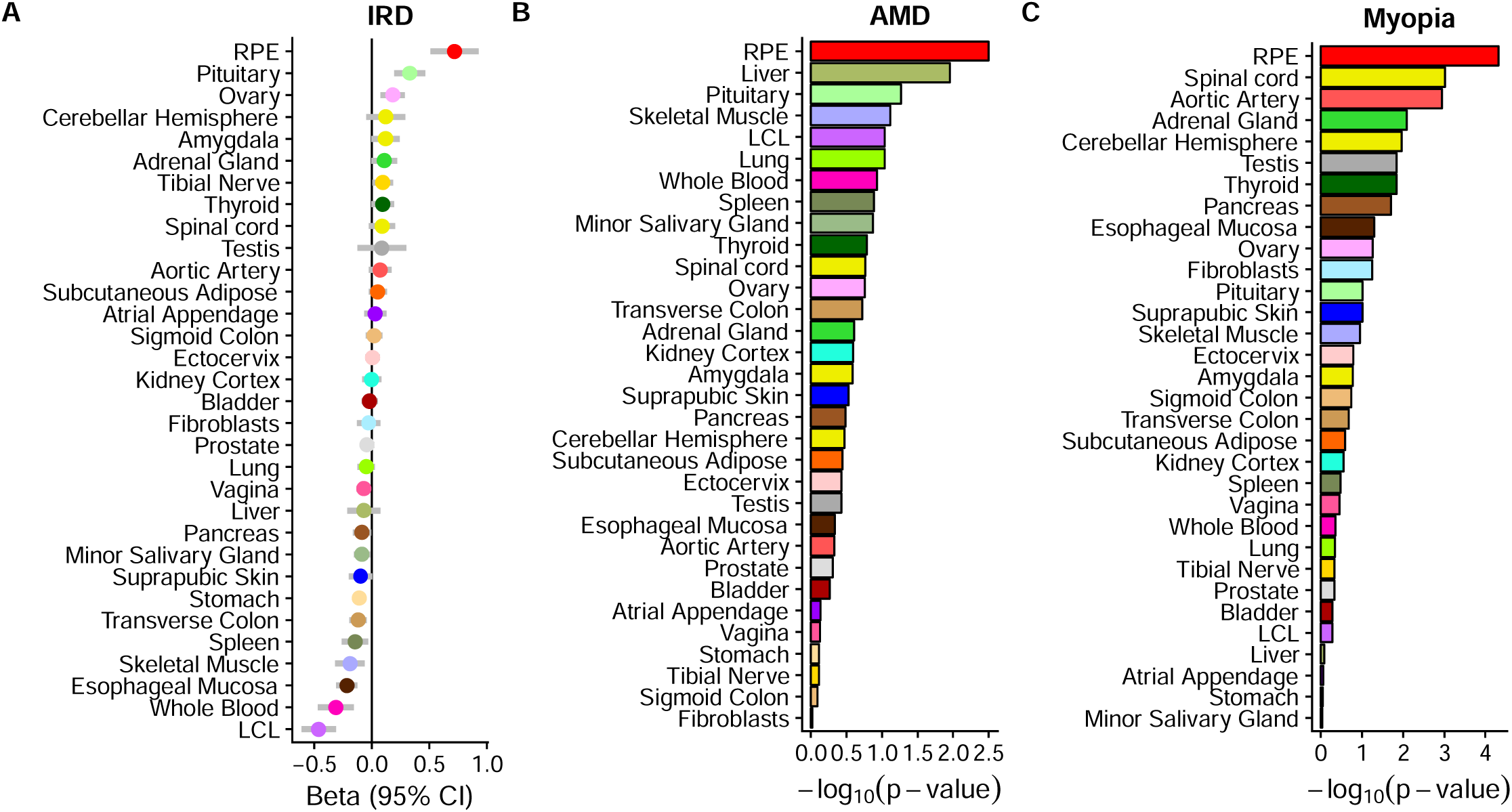
fRPE-selective genes are enriched in monogenic and polygenic diseases. (A) Genes causal for inherited retinal disorders (IRD) have elevated expression in fRPE. (B-C) Variants near RPE-selective genes explain a larger proportion of AMD (B) and myopia (C) risk than those near GTEx tissue-selective genes. The red bar represents fRPE-selective genes with z-score > 4 in both glucose and galactose conditions.

Unlike Mendelian ocular diseases, polygenic ocular disorders are characterized by variants with smaller effect sizes scattered throughout the genome. To determine the heritability explained by fRPE, we performed stratified linkage disequilibrium (LD) score regression using two well-powered GWAS on AMD ^42^ and myopia ^43^. Using a previously established pipeline ^44^, we selected the top 500 tissue-enriched genes and assigned variants within one kilobase of these genes to each tissue (see **Methods**). Risk variants for both complex diseases were better enriched around fRPE-selective genes than GTEx tissue-selective genes (Fig. 3B and C). As an assessment of the robustness of the LD score regression results, we repeated the analysis with the top 200 and 1000 tissue-specific genes. A high ranking for fRPE was consistent across all three cutoffs (Fig. S17).

### Expression and splicing quantitative trait loci discovery

To determine the genetic effects on gene expression in fRPE, we used RASQUAL ^45^ to map eQTL by leveraging both gene-level and allele-specific count information to boost discovery power. Multiple-hypothesis testing for both glucose and galactose conditions was conducted jointly with a hierarchical procedure called TreeQTL ^46^. At FDR < 0.05, we found 726 shared, 272 glucose-specific, and 191 galactose-specific eQTL (Table 1, S10 and S11, Fig. 4A and Fig. S12-13). An example of a glucose-specific eQTL is *ABCA1* (Fig. 4B), which encodes an ATP-binding cassette transporter that regulates cellular cholesterol efflux ^47^. Common variants near *ABCA1* have been associated with glaucoma ^48^ and AMD ^42^. An example of a galactose-specific eQTL is *PRPF8* (Fig. 4C), which encodes a splicing factor ^49^. *PRPF8* mutations are a cause of retinitis pigmentosa ^50^ and lead to RPE dysfunction in a mouse model ^51^. An example of a shared eQTL is *RGR* (Fig. 4D), which encodes a G protein-coupled receptor that is also mutated in retinitis pigmentosa ^52^. We used HOMER ^53^ to identify transcription factor binding motifs enriched around metabolic-specific eQTL (see **Supplementary Methods**). Two motifs, TEAD1 (p < 1×10^−6^) and ZEB1 (p < 1×10^−5^), are amongst the top five motifs in the galactose condition (Table S12). TEAD1 is known to play a role in aerobic glycolysis reprogramming ^54^, and ZEB1 is known to render cells resistant to glucose deprivation ^55^. We did not find enriched motifs in the glucose condition for transcription factors with well-known metabolic functions.

**Figure 4.**
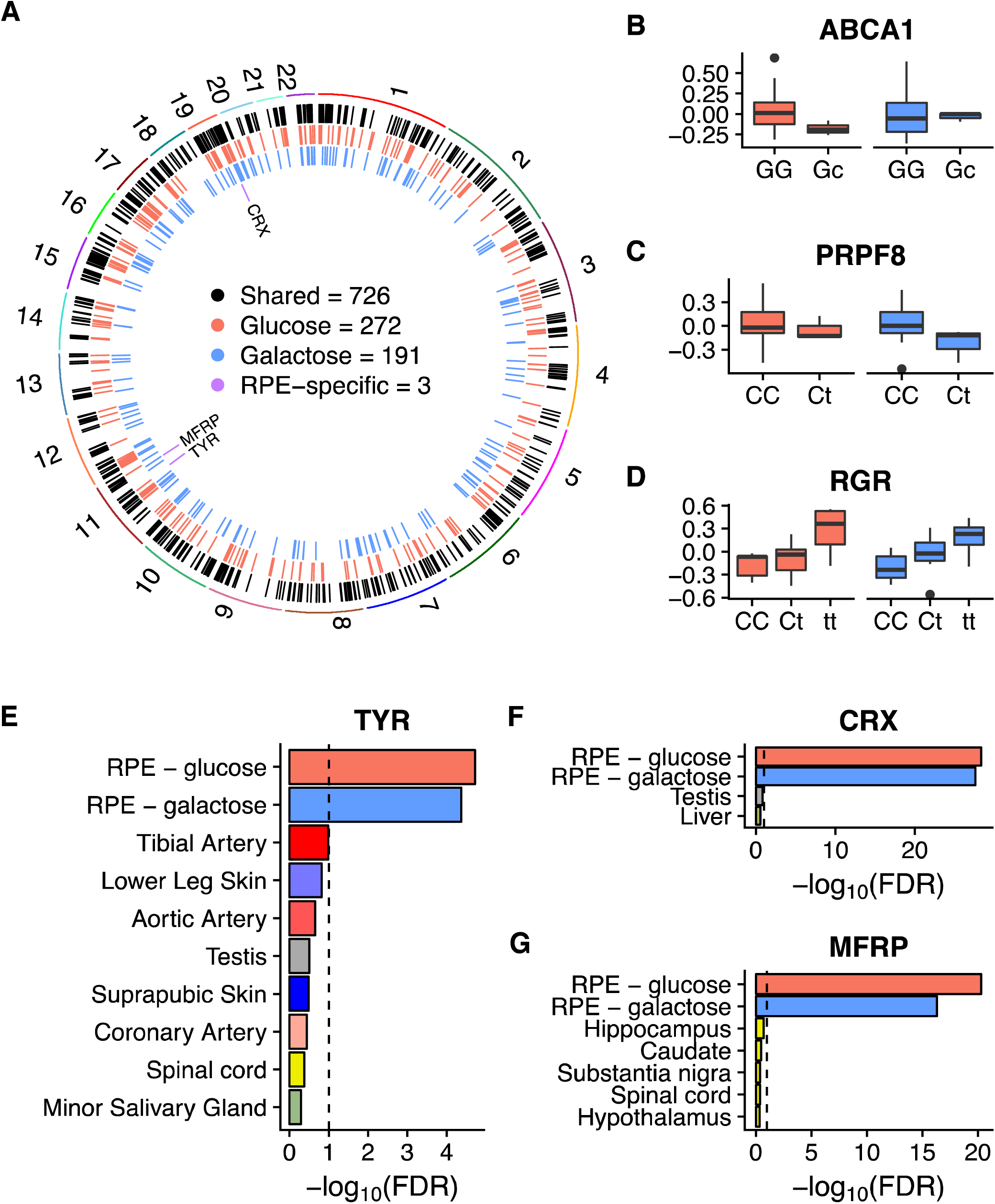
Landscape of genetic regulation of RPE gene expression. (A) We discovered 726, 272, and 191 eQTL that are shared, glucose-specific, and galactose-specific, respectively. Comparison with the GTEx dataset revealed three shared eQTL that are currently unique to fRPE. (B) A glucose-specific eQTL in *ABCA1*. (C) A galactose-specific eQTL in *PRPF8*. (D) A shared eQTL in *RGR*. (E-F) Evidence for fRPE-specificity for three eQTL compared to GTEx. Black dashed lines indicate FDR = 0.1. Minor alleles are indicated by lowercase.

**Table 1.** Expression QTL discoveries

| Trait type | No. tested | No. of eQTL (FDR < 0.05) |  |  |
| --- | --- | --- | --- | --- |
|  |  | Glucose | Galactose | Shared |
| protein coding | 12515 | 262 | 186 | 689 |
| lncRNA | 582 | 10 | 5 | 37 |
| Total | 13097 | 272 | 191 | 726 |

To ascertain whether any genetic regulatory effect is specific to the RPE, we compared fRPE to GTEx eQTL using a previously established two-step FDR approach ^56^. We used fRPE shared eQTL (FDR < 0.05 in both metabolic conditions) as the discovery set to remove any treatment-dependent regulatory effect, and used GTEx eQTL with a relaxed threshold (FDR < 0.1) as the replication set. eQTL from the discovery set not recapitulated in the replicated set were defined as fRPE-selective eQTL. This approach returned three genes (Fig. 4E-G): *TYR*, an oxidase controlling the production of melanin; *CRX*, a transcription factor critical for photoreceptor differentiation; and *MFRP*, a secreted WNT ligand important for eye development. The *TYR* eQTL maps to a variant (rs4547091) previously described as located in an OTX2 binding site and responsible for modulating *TYR* promoter activity in cultured RPE cells ^57^.

We also assessed the genetic effect on splicing by quantifying intron usages with LeafCutter ^58^ and mapping splicing quantitative trait loci (sQTL) with FastQTL ^59^ in permutation mode to obtain intron-level p-values. Following an established approach ^58^, we used a conservative Bonferroni correction across introns within each intron cluster and calculated FDR across cluster-level p-values (see **Supplementary Methods**). We found 210 and 193 sQTL at FDR < 0.05 for glucose and galactose conditions, respectively (Table 2, S13 and S14). The top sQTL in the glucose condition regulates splicing in *ALDH3A2* (FDR < 2.06×10^−9^), which codes for an aldehyde dehydrogenase isozyme involved in lipid metabolism ^60^. Mutations in this gene cause Sjogren-Larsson syndrome ^61^, which can affect the macular RPE ^62^. The top sQTL in the galactose condition regulates splicing of transcripts encoding *CAST*, a calcium-dependent protease inhibitor involved in the turnover of amyloid precursor protein ^63^.

**Table 2.** Splicing QTL discoveries

| Condition | Trait type | No. tested | No. of sQTL |  |  |
| --- | --- | --- | --- | --- | --- |
|  |  |  | FDR < 0.05 | FDR < 0.01 | FDR < 0.001 |
| Glucose | protein coding | 15516 | 205 | 129 | 66 |
|  | lncRNA | 187 | 5 | 4 | 4 |
|  | Total | 15703 | 210 | 133 | 70 |
| Galactose | protein coding | 16239 | 188 | 112 | 56 |
|  | lncRNA | 196 | 5 | 4 | 2 |
|  | Total | 16435 | 193 | 116 | 58 |

### Fine mapping of complex ocular disease risk loci

To assess whether specific instances of GWAS signals can be explained by eQTL or sQTL signals, we performed colocalization analysis with a modified version of eCAVIAR ^64^ (see **Methods**). All variants within a 500-kilobase window around any GWAS (p-value < 1×10^−4^) or QTL (p-value < 1×10^−5^) signal were used as input to eCAVIAR, and any locus with colocalization posterior probability (CLPP) > 0.01 was considered significant. Among four genes exceeding this threshold for eQTL and AMD, *RDH5*, encoding a retinol dehydrogenase that catalyzes the conversion of 11-cis retinol to 11-cis retinal in the visual cycle ^65^, showed the most significant colocalization (Fig. 5A, Table S8). *RHD5* was previously suggested as an AMD candidate gene ^42^, but no mechanism was proposed. Two tightly linked AMD-associated variants (rs3138141 and rs3138142, r^2^=0.98) influence *RDH5* expression (Fig. 5B). The minor haplotype identified by the rs3138141 ‘a’ allele is associated with a significantly smaller percentage of total *RDH5* expression (26.4%) than the major haplotype identified by the ‘C’ allele (73.6%) (Fig. 5G). We found no evidence for an effect on transcripts from the adjacent *BLOC1S1* gene or on *BLOC1S1-RDH5* read-through transcripts. The same haplotype marks an *RDH5* sQTL (Fig. 5A & C) that influences the usage of exon 3 of the transcript; samples that are heterozygous at rs3138141 (Ca) exhibit an average of more than three times the amount of exon 3 skipping compared to CC homozygous samples (Fig. 5H and S22). The same e/sQTL also colocalized with a myopia GWAS signal (Fig. 5D-F, Table S9), suggesting a mechanism for the prior association of the *RDH5* locus with myopia ^66^ and refractive error ^13^. eQTL at *PARP12* and *CLU* also showed evidence of colocalization with AMD and myopia signals, respectively (Fig. 5A & D), but neither locus reached genome-wide significance in the respective GWAS.

**Figure 5.**
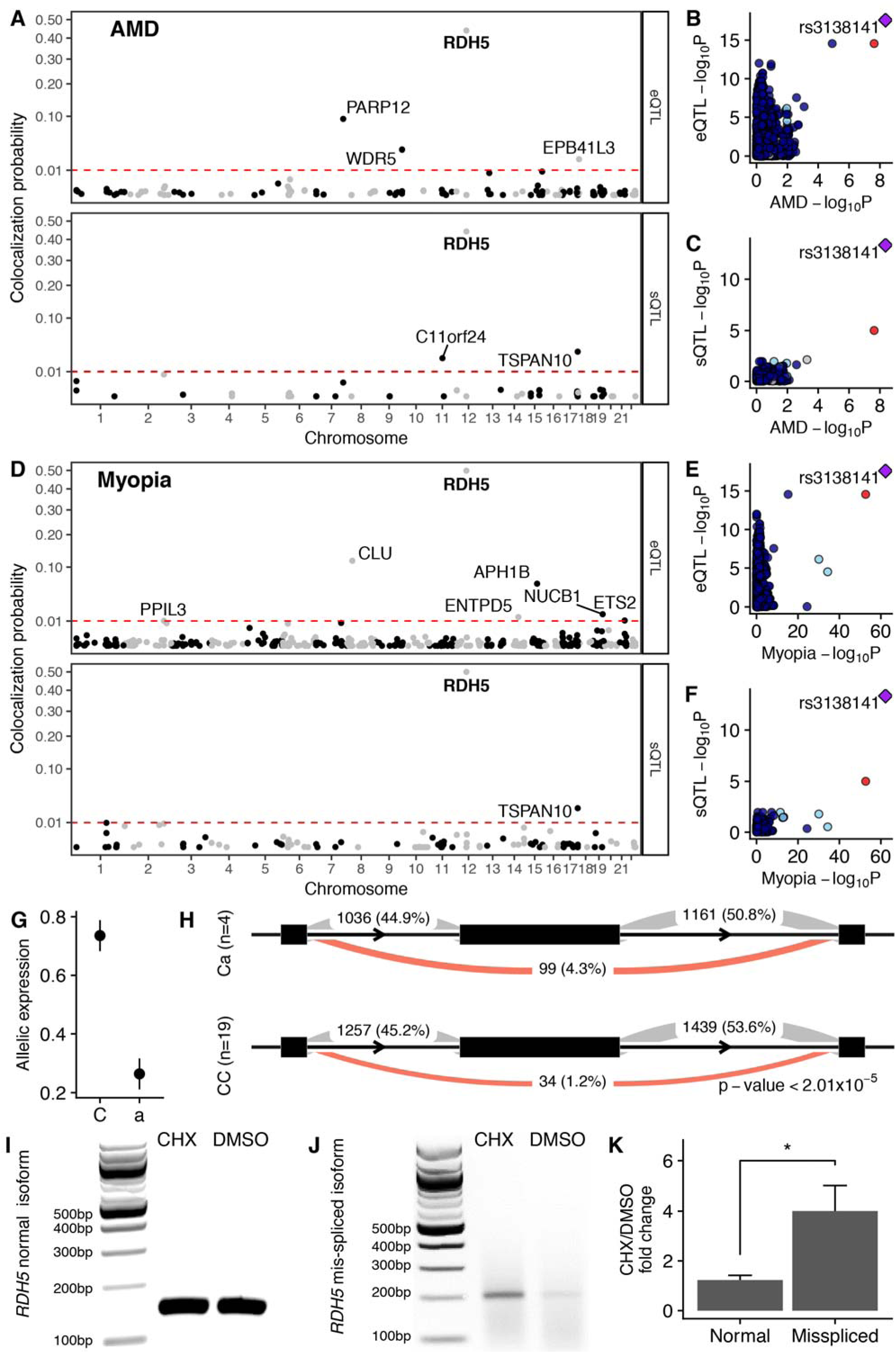
Fine mapping of disease-associated variants using fRPE gene regulation. (A) Colocalization posterior probability for fRPE e/sQTL with AMD. (B-C) Scatter plots demonstrate clear colocalization between AMD GWAS signal at rs3138141 and *RDH5* eQTL (B) and sQTL (C). (D) Colocalization posterior probability for fRPE e/sQTL with myopia. (E-F) Scatter plots demonstrate clear colocalization between myopia GWAS signal at rs3138141, the same variant identified for AMD, and *RDH5* eQTL (E) and sQTL (F). (A-F) Colocalization results are with glucose QTL. Galactose QTL colocalizations can be found in Fig. S18-19. G) Relative allelic expression estimated by RASQUAL with 95% confidence intervals is shown. H) Increased skipping of *RDH5* exon 3 (middle black rectangle) is associated with the minor allele at rs3138141. The average read counts are shown for three splice junctions in groups of fRPE cells with different genotypes. The proportion of counts for all three sites for a given junction and genotype is shown in parenthesis. Exon and intron lengths are not drawn to scale. Minor alleles are indicated by lowercase. I) Gel image showing *RHD5* normal isoform amplified from CHX or DMSO treated ARPE-19 cells. J) Gel image showing *RHD5* mis-spliced isoform amplified from CHX or DMSO treated ARPE-19 cells. K) Relative fold change between CHX and DMSO treatments for normal and mis-spliced RNA isoforms. Error bars indicate standard error of the mean for three independent experiments. *p < 0.05

### Nonsense-mediated decay as a putative mechanism underlying an *RDH*5 eQTL

The association of the rs3138141/2 minor haplotype with both an *RDH5* eQTL and sQTL suggests a mechanistic relationship. The increased skipping of exon 3 (out of 5) associated with the minor haplotype results in more transcripts with a frameshift and a premature termination codon (PTC) near the 5’ end of exon 4. Many mammalian transcripts with PTCs are subject to nonsense-mediated decay (NMD), particularly when the PTC is not located in the last exon ^67^. Treatment of cells with protein synthesis inhibitors such as cycloheximide (CHX) has been shown to increase the abundance of transcripts subject to NMD ^68^. To assess a possible role for NMD in the stability of *RDH5* transcripts, we treated differentiated immortalized human RPE cells (ARPE-19) with CHX and quantified the abundance of the normal and skipped exon 3 isoforms by RTPCR. CHX caused a significant increase in the abundance of the skipped exon 3 isoform as compared to the normal (Fig. 5I-K). These data are consistent with a model in which the minor haplotype promotes the formation of an aberrant *RDH5* mRNA that is subject to NMD, leading to an overall reduction in the steady state levels of *RDH5* transcripts.

## Discussion

The importance of the RPE for development and lifelong homeostasis of the eye has motivated numerous studies of the RPE transcriptome. Several of these studies proposed similar sets of RPE ‘signature’ genes, the largest of which comprises 171 genes ^29-31^. Only 23 of these genes are present among our group of 100 fRPE-selective protein-coding genes. Our approach of comparing fRPE expression levels to GTEx data, which almost exclusively derive from adult autopsy tissue specimens, may have captured genes highly expressed in cultured and/or fetal cells. Absence in GTEx of pure populations of specialized cell types, especially ocular, may explain other genes in our set. Still, many of the genes we identified are known to serve vital functions in the RPE, as demonstrated by pathway enrichment for pigment synthesis and visual processes. We also identified 30 enriched lncRNAs, a class of transcripts not included in previous signature gene sets. The most highly expressed lncRNA in our list, *RMRP*, is critical for proper mitochondrial DNA replication and OXPHOS complex assembly in HeLa cells ^69^, but its role in the RPE has not yet been investigated. RPE-enriched genes whose functions have not been studied in the tissue afford opportunities for advancing understanding of this important epithelial layer.

Our findings have potential implications for phenotypic variability in monogenic ocular diseases. Mutations in all three of the eQTL so far restricted to the fRPE cause monogenic eye diseases. For example, heterozygous mutations in the transcription factor *CRX* cause dominant forms of photoreceptor degeneration, which can exhibit variable age at onset and disease progression among members of the same family ^4^. Genetically encoded variation in the transcript levels of normal or mutant *CRX* alleles may contribute to such variable expressivity. Indeed, mouse models of *CRX*-associated retinopathies provide evidence for a threshold effect in which small changes in expression cause large differences in phenotype ^70^. Mutations in *MRFP* cause extreme hyperopia (farsightedness). Affected individuals usually have two mutant alleles, but inheritance of a lower-expressing normal allele could explain an affected heterozygous individual in a family with otherwise recessive disease ^5^. The substantial number of fRPE eQTL associated with other ocular diseases (Fig. 3A) supports a contribution of common genetic variants to the widespread phenotypic variability observed in monogenic eye disorders.

Our findings also have implications for complex ocular diseases. Evidence suggests that defects in RPE energy metabolism contribute to the pathogenesis of AMD, the hallmark of which is accumulation of cholesterol rich deposits in and around the RPE ^71,72^. Forcing fRPE cells to rely on oxidation of glutamine, the most abundant free amino acid in blood, caused upregulation of genes involved in the synthesis of cholesterol, monounsaturated and polyunsaturated fatty acids, as well as genes associated with lipid import. Transcripts for three of the upregulated genes (*FADS1*, *FADS2*, and *ACAT2*) are increased in macular but not extramacular RPE from individuals with early-stage AMD ^73^.

Co-localization of the same *RDH5* e/sQTL with both AMD and myopia GWAS loci suggests risk mechanisms for these very different complex diseases. The rs3138141/2 minor haplotype confers an elevated risk for AMD ^42^, but is protective for myopia ^13,43,66^. Reduction in *RDH5* activity as a risk factor for AMD is consistent with rare *RDH5* loss-of-function mutations that cause recessive fundus albipunctatus, which can include macular atrophy ^74,75^. More puzzling is the relationship between lower *RDH5* transcript levels (and presumably enzyme activity) and a reduced risk of myopia. RDH5 is best known for its role in the regeneration of 11-cis retinal in the visual cycle, but the enzyme has also been reported to be capable of producing retinoids suitable for retinoic acid signaling ^76,77^. Evidence from animal models implicates retinoic acid in eye growth regulation ^12^, and retinal all-trans retinoic acid levels are elevated in a guinea pig model of myopia ^78^. Thus the same haplotype, which has risen to substantial frequencies in some populations (0.38 minor haplotype frequency in South Asians and 0.19 in Europeans https://www.ncbi.nlm.nih.gov/projects/SNP/), may dampen retinoic acid signaling during eye development and growth, and later contribute to chronic photoreceptor dysfunction in older adults.

The eye is a highly specialized organ with limited representation in large-scale functional genomics datasets. Our analysis of genetic variation and metabolic processes in fRPE cells, even with modest samples sizes, expands our ability to map functional variants with potential to contribute to complex and monogenic eye diseases. Deeper sampling of other ocular cell types, other ethnicities and/or larger sets of samples will likely yield additional discovery of e/sQTL and functional variants involved in genetic eye diseases.

## Methods

### Sample acquisition and cell culture

Primary human fetal RPE (fRPE) lines were isolated by collecting and freezing non-adherent cells cultured in low calcium medium as described ^22^. When needed, fRPE cells were thawed and plated onto 6-well plates in medium as described ^23^ with 15% FBS. The next day, medium was changed to 5% FBS and the cells were allowed to recover for two additional days. Cells were then trypsinized in 0.25% Trypsin-EDTA (Life Technologies Corporation #25200056), resuspended in medium with 15% FBS and plated onto human extracellular matrix-coated (BD Biosciences #354237) Corning 12-well transwells (#3460, Corning Inc., Corning, NY) at 240K cells per transwell. The next day medium was changed to 5% FBS. Cells were cultured for at least 10 weeks to become differentiated (transepithelial resistance of > 200 Ω * cm^2^) and highly pigmented. Medium with 5% FBS was changed every 2-3 days. For the galactose and glucose specific culture conditions, differentiated fRPE cells were cultured for 24 hrs prior to RNA isolation in DMEM medium (#D5030, Sigma) with 1 mM sodium pyruvate (#S8636, Sigma), 4 mM L-glutamine (#25030-081, Life Technologies Corporation), 1% Penicillin-Streptomycin (#15140-122, Life Technologies Corporation), and either 10 mM D(+)-glucose (#G7021, Sigma) or 10 mM D-(+)-galactose (G5388, Sigma) ^28^.

### Genotype data and quality control

All fRPE samples were genotyped on Illumina Infinium Omni2.5-8 BeadChip using the Infinium LCG Assay workflow (https://www.illumina.com/products/by-type/microarray-kits/infinium-omni25-8.html). Samples amplified from 200 ng of genomic DNA were fragmented and hybridized overnight on the Omni2.5-8 BeadChip. The loaded BeadChips went through single-base extension and staining, and were imaged on the iScan machine to obtain genotype information. We used Beagle v4.1 ^24^ to perform genotype imputation and phasing with 1000 Genomes Project Phase 3 reference panel. Prior to imputation and phasing, we filtered the original VCF file to only bi-allelic SNP sites on autosomes and removed sites with more than 5% missing genotypes. Beagle was run with default parameters. After imputation and phasing, we removed variants with allelic r^2^ < 0.8 and removed multi-allelic variants (variants with more than two alleles). Additional quality control considerations are described in **Supplementary Methods**. We compared genotypes between every pair of samples and found a pair of duplicated samples (Fig. S5) and removed one sample arbitrarily.

### Transcriptomic data and quality control

RNA was extracted using TRIzol Reagent (Invitrogen) per manufacturer instructions. RNA sequencing was performed on all samples with an RNA integrity number (RIN) of 8.0 or higher and with at least 500 ng total RNA (Table S4). Stranded, poly-A+ selected RNA-seq libraries were generated using the Illumina TruSeq Stranded mRNA protocol. We performed 75-base paired-end sequencing using an Illumina NextSeq 500. Glucose and galactose samples from each line were sequenced together to minimize batch effects. Raw data was de-multiplexed using bcl2fastq2 from Illumina with default parameters. Reads were aligned against the hg19 human reference genome with STAR (v2.4.2a) ^79^ using GENCODE v19 annotations ^80^ and otherwise default parameters. After alignment, duplicate reads were marked using Picard MarkDuplicates (v2.0.1). We used HTSeq v0.6.0 ^81^ to count the number of reads overlapping each gene based on the GENCODE v19 annotation. We counted reads on the reverse strand (ideal for Illumina’s TruSeq library), required a minimum alignment quality of 10, but otherwise used default parameters. We also quantified RPKM using RNA-SeQC v1.1.8 ^82^ using hg19 reference genome and GENCODE v19 annotation with flags “-noDoC -strictMode” but otherwise default parameters. We quantified allele-specific expression using the createASVCF.sh script from RASQUAL ^45^ with default parameters. For splicing quantification, we used LeafCutter ^58^ to determine intron excision levels with default parameters. We checked the number of uniquely mapped reads to ensure sufficient number of mapped reads. The RNA-seq libraries have a median number of 46.8 million (88.8%) uniquely mapped reads, with an interquartile range of 41.0 to 55.2 million reads (Fig. S6B). We ran VerifyBamID ^83^ with parameters “--ignoreRG --best” on RNA-seq BAM files using genotype VCF files as reference and did not find any sample swaps.

### External datasets

We used GTEx V7 ^18^ as a reference dataset to perform RPE-selective gene and RPE-specific eQTL analyses (see **Supplementary Methods**). The GTEx V7 dataset collected 53 tissues across 714 donors. All tissues across all donors were used in RPE-selective gene analysis. Among the 53 tissues, 48 tissues have sufficient sample size to perform eQTL analysis and were used for RPE-specific eQTL calling. We used two well-powered ocular disorder GWAS datasets to perform colocalization analysis (Table S1). The age-related macular degeneration study ^42^ is a meta-analysis across 26 studies and identified 52 independent GWAS signals, including 16 novel loci. The myopia GWAS was part of a 42-trait GWAS collection aimed at finding shared genetic influences across different traits ^43^. We used ocular disease genes from the Genetic Eye Disease (GEDi) test panel ^41^ (Table S2).

### RPE-selective gene and pathway enrichment analysis

To identify RPE-selective genes (high expression in RPE relative to other tissues), we inferred expression specificity using the following procedure.

1. Calculate the median expression level (x) across all individuals for each tissue.
2. Calculate the mean (μ) and standard deviation (σ) of median expression values across tissues.
3. Derive a z-score for each tissue as follows: z = (x−μ)/σ.
4. Define a gene to be tissue-selective if its z-score is greater than 4.

We filtered out genes on sex and mitochondrial chromosomes, and further filtered out genes in HLA region due to low mappability. To determine whether technical confounders (such as batch effect) affected RPE z-scores, we used a QQ-plot to visualize the z-score of each tissue against the average z-score across tissues. Fig. S9 shows that RPE z-scores situate within the midst of z-scores from GTEx tissues. In fact, the only outlier is testis, which is a known outlier from previous studies. We performed GSEA ^32^ using z-scores as input against GO gene sets from the Molecular Signature Database ^84^ with 10,000 permutations and otherwise default parameters.

### Differential expression and pathway enrichment analysis

We performed differential expression analysis to detect genes whose expression levels were affected by metabolic perturbation. We used DESeq2 ^34^ and corrected for sex, ancestry, RIN, and batch. A simplified mathematical representation is:

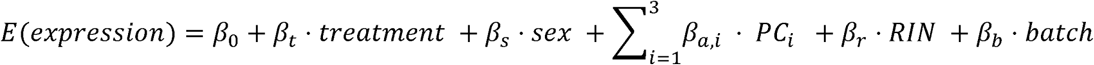

We performed pathway enrichment analysis with GSEA using a ranked gene list with 10,000 permutations but otherwise default parameters. The ranking metric was calculated by multiplying the -log_10_(FDR) by the sign of the effect size from DESeq2. For the pathway database, we used a subset of the Molecular Signatures Database composed of its Hallmark, Biocarta, Reactome, KEGG and GO gene sets ^84^.

### fRPE-selective genes in ocular diseases

We stratified all protein-coding genes into two groups: 1) ocular disease genes (from GEDi, see section **External datasets**) and 2) non-ocular disease genes. To determine whether known ocular disease genes have elevated expression in fRPE, we compared the expression specificity z-score distribution (defined previously) across these two groups with a two-sided t-test. We performed the same analysis for all GTEx tissues as a benchmark. As a control, we repeated this analysis using known epilepsy genes (n = 189) curated from the Invitae epilepsy gene test panel (https://www.invitae.com/en/physician/tests/03401/). GWAS risk loci are frequently enriched around causal genes, which have elevated expression in relevant tissues ^85^. To determine whether variants around RPE-selective genes explain higher disease heritability than expected by chance, we performed stratified LD score regression on tissue-selective genes using a previously established pipeline ^44^. Since LD score regression operates on a variant level, we assigned variants within 1-kb around any exon of tissue-selective genes to each tissue. Although many variants show long-range interaction, we restricted our analysis to a conservative window size to capture only nearby cis-effects. We performed LD score regression on the 200, 500, and 1000 tissue-specific genes (Fig. 3 and Fig. S17).

### Expression QTL mapping and quality control

We extracted hidden factors from RNA sequencing data using surrogate variable analysis (sva) ^86^ separately for glucose and galactose samples. Prior to extracting hidden factors, the raw count gene expression data was library size corrected, variance stabilized, and log2-transformed using the R package DESeq2 ^34^. We ran sva with default parameters and obtained four and five significant surrogate variables for the glucose and galactose conditions, respectively. We performed covariate selection by empirically maximizing the power to detect eQTL. We randomly selected 50 genes from chromosome 22 to perform covariate selection for computational feasibility (RASQUAL ^45^ uses a computationally-intensive iterative fitting procedure) and to avoid overfitting. We added sex, genotype principal components (maximum of three), and surrogate variables sequentially. After multiple hypothesis correction, the number of eAssociations (defined as a SNP-gene pair that passed hierarchical multiple hypothesis testing by TreeQTL ^46^) increased monotonically as the number of covariates, consistent with our intuition that sva only returns significant and independent surrogate variables. Therefore, we decided to use sex, the top three genotype principal components and all surrogate variables (four and five for glucose and galactose conditions, respectively). We mapped eQTL using RASQUAL ^45^ by jointly modeling gene-level and allele-specific read counts. A simplified mathematical representation is as follows:

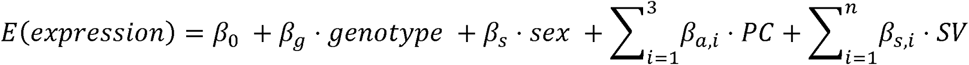

Where n = 4 and 5 for glucose and galactose conditions, respectively. We obtained gene-level and association-level FDR using a hierarchical hypothesis correction procedure implemented in TreeQTL ^46^ using FDR < 0.05 on both gene and association levels. We determined whether the p-values were inflated (e.g. due to model mis-specification) by visualizing their distribution. The distribution suggests that the p-values are slightly conservative (Fig. S10A). As expected, eQTL with low p-values were enriched around transcription start sites (Fig. S10C).

We performed multi-tissue eQTL calling using TreeQTL in multi-tissue calling mode. We set the gene as the first level, the treatment as the second level, and the gene-treatment-SNP as the third level and used the FDR < 0.05 for all three levels. To detect RPE-selective genetic regulatory effects, we compared RPE and GTEx eQTL with a two-step FDR approach as described previously ^56^. In brief, eQTL shared across both conditions in fRPE were selected (FDR<5%). We decided to filter for shared eQTL as they likely reflect regulatory effect not due to treatments. For each eQTL gene, we screened all GTEx tissues for association at a relaxed FDR < 0.1 and defined an eQTL as RPE-selective if no significant association were found in GTEx.

### Splicing QTL calling and quality control

We extracted hidden factors from intron quantification using surrogate variable analysis (sva) ^86^ separately for glucose and galactose samples with default parameters and obtained two significant surrogate variables each for the glucose and galactose conditions. We performed covariate selection by empirically maximizing the power to detect sQTL using intron clusters from chromosome 1 to avoid overfitting. The number of significant sQTL decreased as the number of covariates increased (Fig. S14). This is likely because LeafCutter uses the ratio of read counts between each intron and its intron cluster as the molecular phenotype. Intuitively, if a given batch effect influences the read count of an intron, it will likely influence the quantification of other adjacent introns in the same cluster in the same fashion. Taking the ratio between an intron and its intron cluster effectively cancels out such batch effects.

We mapped sQTL separately for two conditions using FastQTL ^59^ in both nominal and permutation modes and used a simple linear regression:

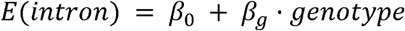

To obtain cluster-level p-values, we used a conservative approach to correct for family-wise error rate with the Bonferroni procedure across introns within each cluster. Global FDR estimates were calculated using the lowest Bonferroni adjusted p-values per cluster. We used FDR < 0.05 as a significance cutoff. As a quality control, we determined whether the p-values were inflated by visualizing their distribution. The p-values showed a uniform distribution with a spike near 0 (Fig. S15A). Further, sQTL with low p-values were enriched around splicing donor and acceptor sites (Fig. S15C), and intronic sQTL SNPs were enriched at intron boundaries (Fig. S15D).

### Fine-mapping of polygenic ocular disease risk loci

We used fRPE eQTL and sQTL information to identify potential causal genes in two well-powered GWAS on age-related macular degeneration and myopia using a modified version of eCAVIAR ^64^. For every significant eQTL, we tested all variants within 500-kb of the lead eQTL SNP for colocalization with GWAS summary statistics. At each candidate locus, we ran FINEMAP ^87^ twice to compute the posterior probability that each individual SNP at the locus was a causal SNP for the GWAS phenotype and fRPE e/sQTL. We then processed the FINEMAP results to compute a colocalization posterior probability (CLPP) using the method described by eCAVIAR ^64^. We defined any locus with CLPP > 0.01 to have sufficient evidence for colocalization. At loci that showed colocalization between RPE eQTL and GWAS associations, we performed the colocalization tests again using eQTL from each of 44 GTEx tissues. To determine whether any potential causal genes act primarily through fRPE, we repeated colocalization analysis with GTEx eQTL.

### Experimental validation

ARPE-19 cells were differentiated for 3 months in 6-well plates (Corning) in medium containing 3 mM pyruvate ^88^ and treated with 100 μg/mL cycloheximide (CHX; Sigma) or vehicle (DMSO) for 3 hr. Cells were then collected, RNA was extracted by TRIzol (Invitrogen), and cDNA was synthesized with an iScript™ cDNA Synthesis Kit (Bio-RAD). Oligonucleotide primers were designed to specifically amplify the normal or misspliced isoforms of the *RDH5* transcript. For the normal isoform, the forward primer (ggggctactgtgtctccaaa) was located in exon 3 and the reverse primer (tgcagggttttctccagact) was located in exon 4, with an expected product size of 151 bp. The amplification conditions were: 94°C 2 min followed by 38 cycles of 94°C 30 sec, 60°C 30 sec, 72°C 15 sec. For the mis-spliced isoform, the forward primer (gatgcacgttaaggaagcag/gcg) spanned the exon 2/4 junction with the three bases at the 3’ end located in exon 4. The reverse primer (gcgctgttgcattttcaggt) was located in exon 5. The expected product size is 204 bp. The amplification conditions were: 94°C 2 min followed by 50 cycles of 94°C 30 sec, 60°C 30 sec, 72°C 15 sec. AmpliTaq (ThermoFisher) and 2.5 mM MgCl2 were used for all reactions. The identities of the normal and mis-spliced PCR products were confirmed by Sanger sequencing. For quantification, PCR products were resolved on 2% agarose gels containing ethidium bromide and imaged using a Bio-Rad ChemiDoc Touch Imaging System. Equal-sized boxes were drawn around bands for the CHX and DMSO samples, grayscale values were measured by ImageJ (NIH), and the relative fold change was calculated (mean ± SEM; three independent experiments). A one-sided Students *t*-test was used to assess the statistical significance of a model under which CHX increased product abundance.

## Data Availability

All relevant data are available in the **Supplementary Data** files. Full eQTL and sQTL summary statistics have been made deposited into Box: https://stanford.box.com/s/asrxy0o66xxe1j7mfj56p3z3d405gijj and are available at http://montgomerylab.stanford.edu/resources.html.

## Code Availability

Code to reproduce all analyses in this manuscript has been deposited on GitHub: https://github.com/boxiangliu/rpe

## Acknowledgements

This work was supported by a Stanford Center for Computational, Evolutionary and Human Genomics Predoctoral Fellowship and the National Key R&D Program of China (2016YFD0400800) (to BL); T32EY20485 (to MAC); the Edward Mallinckrodt Jr. Foundation and NIH grants R33HL120757, U01HG009431, R01MH101814, and R01HG008150 (to SBM); The Macular Degeneration Research Program of the BrightFocus Foundation, the Foundation Fighting Blindness, and R01EY025790 (to DV); and P30EY026877. DB is the Dolly Green Professor of Ophthalmology at UCLA. The authors thank 23andMe employees and participants for providing summary statistics for the myopia GWAS.

## Author Contributions

Conceived and designed experiments: MAC, SBM, DV. Provided critical reagents and expertise: DB, JH. Performed experiments: GB, MAC, MC, BL. Analyzed data: BB, NSB, MJG, BL, XL, SBM, DV. Wrote the paper: BB, MAC, BL, SBM, DV.

## References

1 McKusick, V. A. Mendelian Inheritance in Man and its online version, OMIM. The American Journal of Human Genetics 80, 588–604, doi:10.1086/514346 (2007).

2 Boon, C. J. F. et al. The spectrum of retinal dystrophies caused by mutations in the peripherin/RDS gene. Progress in Retinal and Eye Research 27, 213–235, doi:10.1016/j.preteyeres.2008.01.002 (2008).

3 Nash, B. M., Wright, D. C., Grigg, J. R., Bennetts, B. & Jamieson, R. V. Retinal dystrophies, genomic applications in diagnosis and prospects for therapy. Translational Pediatrics 4, 139–163, doi:10.3978/j.issn.2224-4336.2015.04.03 (2015).

4 Paunescu, K., Preising, M. N., Janke, B., Wissinger, B. & Lorenz, B. Genotype–Phenotype Correlation in a German Family with a Novel Complex CRX Mutation Extending the Open Reading Frame. Ophthalmology 114, 1348–1357.e1341, doi:10.1016/j.ophtha.2006.10.034 (2007).

5 Sundin, O. H. et al. Extreme hyperopia is the result of null mutations in MFRP, which encodes a Frizzled-related protein. Proceedings of the National Academy of Sciences 102, 9553–9558, doi:10.1073/pnas.0501451102 (2005).

6 Vaclavik, V., Gaillard, M. C., Tiab, L., Schorderet, D. F. & Munier, F. L. Variable phenotypic expressivity in a Swiss family with autosomal dominant retinitis pigmentosa due to a T494M mutation in the PRPF3 gene. Molecular Vision 16, 467–475 (2010).

7 Sergouniotis, P. I. et al. Phenotypic Variability in RDH5 Retinopathy (Fundus Albipunctatus). Ophthalmology 118, 1661–1670, doi:10.1016/j.ophtha.2010.12.031 (2011).

8 Llavona, P. et al. Allelic Expression Imbalance in the Human Retinal Transcriptome and Potential Impact on Inherited Retinal Diseases. Genes 8, 283, doi:10.3390/genes8100283 (2017).

9 MacArthur, J. et al. The new NHGRI-EBI Catalog of published genome-wide association studies (GWAS Catalog). Nucleic Acids Research 45, D896–D901, doi:10.1093/nar/gkw1133 (2017).

10 Bressler, N. M. Age-Related Macular Degeneration Is the Leading Cause of Blindness…JAMA 291, 1900–1901, doi:10.1001/jama.291.15.1900 (2004).

11 Swaroop, A., Chew, E. Y., Bowes Rickman, C. & Abecasis, G. R. Unraveling a multifactorial late-onset disease: from genetic susceptibility to disease mechanisms for age-related macular degeneration. Annual Review of Genomics and Human Genetics 10, 19–43, doi:10.1146/annurev.genom.9.081307.164350 (2009).

12 Zhang, Y. & Wildsoet, C. F. RPE and Choroid Mechanisms Underlying Ocular Growth and Myopia. Progress in Molecular Biology and Translational Science 134, 221–240, doi:10.1016/bs.pmbts.2015.06.014 (2015).

13 Tedja, M. S. et al. Genome-wide association meta-analysis highlights light-induced signaling as a driver for refractive error. Nature Genetics 50, 834–848, doi:10.1038/s41588-018-0127-7 (2018).

14 Holden, B. A. et al. Global Prevalence of Myopia and High Myopia and Temporal Trends from 2000 through 2050. Ophthalmology 123, 1036–1042, doi:10.1016/j.ophtha.2016.01.006 (2016).

15 Nicolae, D. L. et al. Trait-Associated SNPs Are More Likely to Be eQTLs: Annotation to Enhance Discovery from GWAS. PLoS genetics 6, e1000888, doi:10.1371/journal.pgen.1000888 (2010).

16 Gusev, A. et al. Partitioning Heritability of Regulatory and Cell-Type-Specific Variants across 11 Common Diseases. The American Journal of Human Genetics 95, 535–552, doi:10.1016/j.ajhg.2014.10.004 (2014).

17 Nica, A. C. & Dermitzakis, E. T. Expression quantitative trait loci: present and future. Philosophical Transactions of the Royal Society B: Biological Sciences 368, 20120362–20120362, doi:10.1098/rstb.2012.0362 (2013).

18 GTEx Consortium et al. Genetic effects on gene expression across human tissues. Nature 550, 204-213, doi:10.1038/nature24277 (2017).

19 Raymond, S. M. & Jackson, I. J. The retinal pigmented epithelium is required for development and maintenance of the mouse neural retina. Current Biology 5, 1286–1295, doi:10.1016/S0960-9822(95)00255-7 (1995).

20 Strauss, O. The Retinal Pigment Epithelium in Visual Function. Physiological Reviews 85, 845–881, doi:10.1152/physrev.00021.2004 (2005).

21 Vollrath, D. et al. Tyro3 Modulates Mertk-Associated Retinal Degeneration. PLoS genetics 11, e1005723, doi:10.1371/journal.pgen.1005723 (2015).

22 Hu, J. & Bok, D. Culture of highly differentiated human retinal pigment epithelium for analysis of the polarized uptake, processing, and secretion of retinoids. Methods in Molecular Biology 652, 55–73, doi:10.1007/978-1-60327-325-1_2 (2010).

23 Maminishkis, A. et al. Confluent Monolayers of Cultured Human Fetal Retinal Pigment Epithelium Exhibit Morphology and Physiology of Native Tissue. Investigative Ophthalmology & Visual Science 47, 3612–3624, doi:10.1167/iovs.05-1622 (2006).

24 Browning, B. L. & Browning, S. R. Genotype Imputation with Millions of Reference Samples. The American Journal of Human Genetics 98, 116–126, doi:10.1016/j.ajhg.2015.11.020 (2016).

25 1000 Genomes Project Consortium et al. A global reference for human genetic variation. Nature 526, 68–74, doi:10.1038/nature15393 (2015).

26 Folmes, C. D. L., Dzeja, P. P., Nelson, T. J. & Terzic, A. Metabolic plasticity in stem cell homeostasis and differentiation. Cell Stem Cell 11, 596–606, doi:10.1016/j.stem.2012.10.002 (2012).

27 Terluk, M. R. et al. Investigating mitochondria as a target for treating age-related macular degeneration. Journal of Neuroscience 35, 7304–7311, doi:10.1523/JNEUROSCI.0190-15.2015 (2015).

28 Gohil, V. M. et al. Nutrient-sensitized screening for drugs that shift energy metabolism from mitochondrial respiration to glycolysis. Nature Biotechnology 28, 249–255, doi:10.1038/nbt.1606 (2010).

29 Bennis, A. et al. Comparison of Mouse and Human Retinal Pigment Epithelium Gene Expression Profiles: Potential Implications for Age-Related Macular Degeneration. PloS one 10, e0141597, doi:10.1371/journal.pone.0141597 (2015).

30 Liao, J.-L. et al. Molecular signature of primary retinal pigment epithelium and stem-cell-derived RPE cells. Human Molecular Genetics 19, 4229–4238, doi:10.1093/hmg/ddq341 (2010).

31 Strunnikova, N. V. et al. Transcriptome analysis and molecular signature of human retinal pigment epithelium. Human Molecular Genetics 19, 2468–2486, doi:10.1093/hmg/ddq129 (2010).

32 Subramanian, A. et al. Gene set enrichment analysis: A knowledge-based approach for interpreting genome-wide expression profiles. Proceedings of the National Academy of Sciences 102, 15545–15550, doi:10.1073/pnas.0506580102 (2005).

33 Ashburner, M. et al. Gene Ontology: tool for the unification of biology. Nature Genetics 25, 25–29, doi:10.1038/75556 (2000).

34 Love, M. I., Huber, W. & Anders, S. Moderated estimation of fold change and dispersion for RNA-seq data with DESeq2. Genome Biology 15, 550, doi:10.1186/s13059-014-0550-8 (2014).

35 Paton, C. M. & Ntambi, J. M. Biochemical and physiological function of stearoyl-CoA desaturase. American Journal of Physiology - Endocrinology and Metabolism 297, E28–E37, doi:10.1152/ajpendo.90897.2008 (2009).

36 Samuel, W. et al. Regulation of stearoyl coenzyme A desaturase expression in human retinal pigment epithelial cells by retinoic acid. The Journal of Biological Chemistry 276, 28744–28750, doi:10.1074/jbc.M103587200 (2001).

37 Yang, T. et al. Crucial Step in Cholesterol Homeostasis: Sterols Promote Binding of SCAP to INSIG-1, a Membrane Protein that Facilitates Retention of SREBPs in ER. Cell 110, 489–500, doi:10.1016/S0092-8674(02)00872-3 (2002).

38 Aledo, R. et al. Genetic basis of mitochondrial HMG-CoA synthase deficiency. Human Genetics 109, 19–23 (2001).

39 Reyes-Reveles, J. et al. Phagocytosis-dependent ketogenesis in retinal pigment epithelium. The Journal of Biological Chemistry 292, 8038–8047, doi:10.1074/jbc.M116.770784 (2017).

40 Slowikowski, K., Hu, X. & Raychaudhuri, S. SNPsea: an algorithm to identify cell types, tissues and pathways affected by risk loci. Bioinformatics 30, 2496–2497, doi:10.1093/bioinformatics/btu326 (2014).

41 Consugar, M. B. et al. Panel-based genetic diagnostic testing for inherited eye diseases is highly accurate and reproducible, and more sensitive for variant detection, than exome sequencing. Genetics in Medicine 17, 253–261, doi:10.1038/gim.2014.172 (2015).

42 Fritsche, L. G. et al. A large genome-wide association study of age-related macular degeneration highlights contributions of rare and common variants. Nature Genetics 48, 134–143, doi:10.1038/ng.3448 (2015).

43 Pickrell, J. K. et al. Detection and interpretation of shared genetic influences on 42 human traits. Nature Genetics 48, 709–717, doi:10.1038/ng.3570 (2016).

44 Boyle, E. A., Li, Y. I. & Pritchard, J. K. An Expanded View of Complex Traits: From Polygenic to Omnigenic. Cell 169, 1177–1186, doi:10.1016/j.cell.2017.05.038 (2017).

45 Kumasaka, N., Knights, A. J. & Gaffney, D. J. Fine-mapping cellular QTLs with RASQUAL and ATAC-seq. Nature Genetics 48, 206–213, doi:10.1038/ng.3467 (2016).

46 Peterson, C. B., Bogomolov, M., Benjamini, Y. & Sabatti, C. TreeQTL: hierarchical error control for eQTL findings. Bioinformatics 32, 2556–2558, doi:10.1093/bioinformatics/btw198 (2016).

47 Schmitz, G. & Langmann, T. Structure, function and regulation of the ABC1 gene product. Current Opinion in Lipidology 12, 129 (2001).

48 Chen, Y. et al. Common variants near *ABCA1* and in *PMM2* are associated with primary open-angle glaucoma. Nature Genetics 46, 1115–1119, doi:10.1038/ng.3078 (2014).

49 Luo, H. R., Moreau, G. A., Levin, N. & Moore, M. J. The human Prp8 protein is a component of both U2- and U12-dependent spliceosomes. RNA 5, 893–908 (1999).

50 Tanackovic, G. et al. PRPF mutations are associated with generalized defects in spliceosome formation and pre-mRNA splicing in patients with retinitis pigmentosa. Human Molecular Genetics 20, 2116–2130, doi:10.1093/hmg/ddr094 (2011).

51 Farkas, M. H. et al. Mutations in pre-mRNA processing factors 3, 8, and 31 cause dysfunction of the retinal pigment epithelium. The American Journal of Pathology 184, 2641–2652, doi:10.1016/j.ajpath.2014.06.026 (2014).

52 Morimura, H., Saindelle-Ribeaudeau, F., Berson, E. L. & Dryja, T. P. Mutations in RGR, encoding a light-sensitive opsin homologue, in patients with retinitis pigmentosa. Nature Genetics 23, 393–394, doi:10.1038/70496 (1999).

53 Heinz, S. et al. Simple Combinations of Lineage-Determining Transcription Factors Prime cis-Regulatory Elements Required for Macrophage and B Cell Identities. Molecular Cell 38, 576–589, doi:10.1016/j.molcel.2010.05.004 (2010).

54 Enzo, E. et al. Aerobic glycolysis tunes YAP/TAZ transcriptional activity. Embo J 34, 1349–1370, doi:10.15252/embj.201490379 (2015).

55 Kanska, J. et al. Glucose deprivation elicits phenotypic plasticity via ZEB1-mediated expression of NNMT. Oncotarget 8, 26200–26220, doi:10.18632/oncotarget.15429 (2017).

56 Barreiro, L. B. et al. Deciphering the genetic architecture of variation in the immune response to Mycobacterium tuberculosis infection. Proceedings of the National Academy of Sciences of the United States of America 109, 1204–1209, doi:10.1073/pnas.1115761109 (2012).

57 Reinisalo, M., Putula, J., Mannermaa, E., Urtti, A. & Honkakoski, P. Regulation of the human tyrosinase gene in retinal pigment epithelium cells: the significance of transcription factor orthodenticle homeobox 2 and its polymorphic binding site. Molecular Vision 18, 38–54 (2012).

58 Li, Y. I. et al. Annotation-free quantification of RNA splicing using LeafCutter. Nature Genetics 50, 151-158, doi:10.1038/s41588-017-0004-9 (2018).

59 Ongen, H., Buil, A., Brown, A. A., Dermitzakis, E. T. & Delaneau, O. Fast and efficient QTL mapper for thousands of molecular phenotypes. Bioinformatics 32, 1479–1485, doi:10.1093/bioinformatics/btv722 (2016).

60 Kelson, T. L., Secor McVoy, J. R. & Rizzo, W. B. Human liver fatty aldehyde dehydrogenase: microsomal localization, purification, and biochemical characterization. Biochimica et Biophysica Acta 1335, 99–110, doi:10.1016/S0304-4165(96)00126-2 (1997).

61 Nakahara, K. et al. The Sjögren-Larsson Syndrome Gene Encodes a Hexadecenal Dehydrogenase of the Sphingosine 1-Phosphate Degradation Pathway. Molecular Cell 46, 461–471, doi:10.1016/j.molcel.2012.04.033 (2012).

62 Nilsson, S. E. & Jagell, S. Lipofuscin and melanin content of the retinal pigment epithelium in a case of Sjögren-Larsson syndrome. The British Journal of Ophthalmology 71, 224–226, doi:10.1136/bjo.71.3.224 (1987).

63 Hanna, R. A., Campbell, R. L. & Davies, P. L. Calcium-bound structure of calpain and its mechanism of inhibition by calpastatin. Nature 456, 409–412, doi:10.1038/nature07451 (2008).

64 Hormozdiari, F. et al. Colocalization of GWAS and eQTL Signals Detects Target Genes. The American Journal of Human Genetics 99, 1245–1260, doi:10.1016/j.ajhg.2016.10.003 (2016).

65 Sahu, B. & Maeda, A. Retinol Dehydrogenases Regulate Vitamin A Metabolism for Visual Function. Nutrients 8, 746, doi:10.3390/nu8110746 (2016).

66 Kiefer, A. K. et al. Genome-Wide Analysis Points to Roles for Extracellular Matrix Remodeling, the Visual Cycle, and Neuronal Development in Myopia. PLoS genetics 9, e1003299, doi:10.1371/journal.pgen.1003299 (2013).

67 Nickless, A., Bailis, J. M. & You, Z. Control of gene expression through the nonsense-mediated RNA decay pathway. Cell & Bioscience 7, 26, doi:10.1186/s13578-017-0153-7 (2017).

68 Carter, M. S. et al. A regulatory mechanism that detects premature nonsense codons in T-cell receptor transcripts in vivo is reversed by protein synthesis inhibitors in vitro. The Journal of Biological Chemistry 270, 28995–29003 (1995).

69 Noh, J. H. et al. HuR and GRSF1 modulate the nuclear export and mitochondrial localization of the lncRNA RMRP. Genes & Development 30, 1224–1239, doi:10.1101/gad.276022.115 (2016).

70 Ruzycki, P. A., Tran, N. M., Kolesnikov, A. V., Kefalov, V. J. & Chen, S. Graded gene expression changes determine phenotype severity in mouse models of CRX-associated retinopathies. Genome Biology 16, 114, doi:10.1186/s13059-015-0732-z (2015).

71 Curcio, C. A. et al. Esterified and unesterified cholesterol in drusen and basal deposits of eyes with age-related maculopathy. Experimental Eye Research 81, 731–741 (2005).

72 Pikuleva, I. A. & Curcio, C. A. Cholesterol in the retina: The best is yet to come. Progress in Retinal and Eye Research 41, 64–89 (2014).

73 Ashikawa, Y. et al. Potential protective function of the sterol regulatory element binding factor 1-fatty acid desaturase 1/2 axis in early-stage age-related macular degeneration. Heliyon 3, e00266, doi:10.1016/j.heliyon.2017.e00266 (2017).

74 Yamamoto, H. et al. Mutations in the gene encoding 11-cis retinol dehydrogenase cause delayed dark adaptation and fundus albipunctatus. Nature Genetics 22, 188–191 (1999).

75 Yamamoto, H. et al. A novel RDH5 gene mutation in a patient with fundus albipunctatus presenting with macular atrophy and fading white dots. American Journal of Ophthalmology 136, 572–574 (2003).

76 Duester, G. Families of retinoid dehydrogenases regulating vitamin A function. European Journal of Biochemistry 267, 4315–4324, doi:10.1046/j.1432-1327.2000.01497.x (2001).

77 Nadauld, L. D. et al. Dual roles for adenomatous polyposis coli in regulating retinoic acid biosynthesis and Wnt during ocular development. Proceedings of the National Academy of Sciences 103, 13409-13414, doi:10.1073/pnas.0601634103 (2006).

78 McFadden, S. A., Howlett, M. H. C. & Mertz, J. R. Retinoic acid signals the direction of ocular elongation in the guinea pig eye. Vision Research 44, 643–653, doi:10.1016/j.visres.2003.11.002 (2004).

79 Dobin, A. et al. STAR: ultrafast universal RNA-seq aligner. Bioinformatics 29, bts635-621, doi:10.1093/bioinformatics/bts635 (2012).

80 Harrow, J. et al. GENCODE: the reference human genome annotation for The ENCODE Project. Genome research 22, 1760–1774, doi:10.1101/gr.135350.111 (2012).

81 Anders, S., Pyl, P. T. & Huber, W. HTSeq--a Python framework to work with high-throughput sequencing data. Bioinformatics 31, 166–169, doi:10.1093/bioinformatics/btu638 (2015).

82 Deluca, D. S. et al. RNA-SeQC: RNA-seq metrics for quality control and process optimization. Bioinformatics 28, 1530–1532, doi:10.1093/bioinformatics/bts196 (2012).

83 Jun, G. et al. Detecting and estimating contamination of human DNA samples in sequencing and array-based genotype data. The American Journal of Human Genetics 91, 839–848, doi:10.1016/j.ajhg.2012.09.004 (2012).

84 Liberzon, A. et al. Molecular signatures database (MSigDB) 3.0. Bioinformatics 27, 1739–1740, doi:10.1093/bioinformatics/btr260 (2011).

85 Finucane, H. K. et al. Heritability enrichment of specifically expressed genes identifies disease-relevant tissues and cell types. Nature Genetics 50, 621–629, doi:10.1038/s41588-018-0081-4 (2018).

86 Leek, J. T. & Storey, J. D. Capturing heterogeneity in gene expression studies by surrogate variable analysis. PLoS genetics 3, 1724–1735, doi:10.1371/journal.pgen.0030161 (2007).

87 Benner, C. et al. FINEMAP: efficient variable selection using summary data from genome-wide association studies. Bioinformatics 32, 1493–1501, doi:10.1093/bioinformatics/btw018 (2016).

88 Samuel, W. et al. Appropriately differentiated ARPE-19 cells regain phenotype and gene expression profiles similar to those of native RPE cells. Molecular Vision 23, 60–89 (2017).

